# Immune state networks of wild and laboratory mice

**DOI:** 10.1101/638445

**Authors:** Elohim Fonseca dos Reis, Mark Viney, Naoki Masuda

## Abstract

The mammalian immune system protects individuals from infection and disease. It is a complex system of interacting cells and molecules and extensive work, principally with laboratory mice, has investigated its function. Wild and laboratory animals lead very different lives, and this is reflected in there being substantial immunological differences between them. Here we use network analyses to study a unique data set of 120 immune measures of wild and laboratory mice, where immune measures define nodes and correlations of immune measures across individual mice define edges between immune measures. To date, there has only been very limited network analyses of the immune system, which is surprising because such analyses may be important to better understand its organisation and functionality. We found that the immunological networks of wild and laboratory mice were similar in some aspects of their mesoscale structure, particularly concerning cytokine response communities. However, we also identified notable differences in node membership of network communities between the wild and laboratory networks, pointing to how the same immune system acts and interacts differently in wild and in laboratory mice. These results show the utility of network analysis in understanding immune responses and also the importance of studying wild animals in additional to laboratory animals.

**Author summary:** The mammalian immune system is a complex system that protects individuals from infection and disease. Most of our understanding of the immune system comes from studies of laboratory animals, particularly mice. However, wild and laboratory animals lead very different lives, potentially leading to substantial immunological differences between them and so possibly limiting the utility of laboratory animals as informative model systems. As a complex interacting set of cells and molecules, the immune system is a biological network. Therefore, we used network analyses to study the immune system, specifically a unique data set of immune measures of wild and laboratory mice, where 120 different immune measures define nodes of the network. We found that the networks of wild and laboratory mice were similar in some aspects of their grouping structure, particularly concerning communities of nodes of cytokine responses. However, we also identified notable differences in node membership of communities between the wild and laboratory networks, pointing to how the same immune system behaves differently in wild and in laboratory mice. These results show the utility of network analysis in understanding immune responses and also the importance of studying wild animals in addition to laboratory animals.

## Introduction

The vertebrate immune system defends animals against infection and disease. Understanding how the immune system functions has been a long-standing area of study, not only to understand the basic biology of this important system, but also to be able to manipulate it for therapeutic benefit. Almost all that we know about the mammalian immune system has been discovered by studying laboratory animals, mainly mice. Remarkably little is known about the immune responses of wild mammals, nor how well the immune responses of laboratory animals reflect those of wild animals. The operation of the immune system is affected by the environment – especially infections – and wild animals are continuously exposed to a myriad of infections stimulating immune responses, but in contrast, laboratory animals do not have this high-level exposure. Indeed, the few studies of the immune state of wild animals have shown quantitative differences in measures of immune state between wild and laboratory animals [1].

The vertebrate immune system is a complex of interacting cells and soluble molecules whose function, at its heart, is to recognise foreign antigen molecules, to then remove, destroy or nullify them, while usually retaining a molecular memory of the antigen in question. The multiple components of the vertebrate immune system are typically categorised into cellular and humoral components, both of which contribute to the innate and adaptive parts of the immune response. The immune system is dynamic, so that different components of the immune system respond depending on the type of antigen and its location in the individual. For example, immune responses to viruses infecting lung cells are qualitatively very different to the immune responses directed against macroscopic worms living in the gut [2]. Pathogens are not passive partners in the face of the immune responses directed against them, and so combat the immune response via a number of strategies. These strategies include molecularly hiding from the immune system, actively immunomodulating the immune system for their own benefit, or by changing their antigenic repertoire, so staying-ahead of an immunological memory already acquired against them [3]. Effector cells and molecules of the immune system act against invading pathogens, but these effects can also cause harm to the individual, so-called immunopathology [4]. Therefore, a tightly-controlled, well regulated immune system is needed to protect an individual from pathogen-induced harm, without causing direct harm to the individual in the process.

The immune system is a regulated, homeostatic system, typically responding to multiple antigenic stimulations at once. The principal actors of the immune system are populations of cells, including T cells (including CD4^+^ T helper cells that facilitate immune function and CD8^+^ T cytotoxic cells that act against infected cells), B cells (that are responsible for antibody responses), as well as other populations of cells including Natural Killer cells, Dendritic cells, and other myeloid cells. Critical to the successful function of the immune system is regulation and coordination of the components of the immune system, which is achieved by a suite of soluble signalling molecules called cytokines. Cytokines include interferons (molecules produced by host cells upon infection) and interleukins (cytokines produced by white blood cells, leukocytes) among others. Cytokines allow cells of the immune system to signal their state, and so provide positive and negative regulation of different components of the immune system. Immune systems that are in different functional and effector states will therefore likely have different underlying quantitative networks of the coordinating cytokine molecules.

Because it is a complex interacting set of cells and molecules, the immune system is a biological network. Applying network analysis to immunological data may therefore provide new insights into the function and organisation of the immune system. Because of the different immunological state of wild and laboratory animals [1, 5], their immune systems may also differ in the structure of interactions among different immune components, which will be revealed by network analysis.

Here we apply network analysis to a data set that has comprehensively measured the immune state of wild mice and laboratory mice, affording us the opportunity to examine the commonalities and the differences between the behaviour of the immune system in these animals. In particular, we analyse the community structure [6], which aims to reveal groups (i.e., communities) of densely connected nodes (which here are measures of the immune response) and the connectivity among the detected communities. We then compare the wild and laboratory mice in terms of community structure and also compare the biological groupings of immune measures and the grouping derived from community detection. As such, this analysis provides a comprehensive network analysis of the immune system, and gives new insights into the functional immunological differences between wild and laboratory animals.

## Methods

### Data

We used a data set of the immune state of 460 wild mice (*Mus musculus domesticus*), collected from 12 sites in the southern UK, and 102 laboratory mice. We assembled this data set in previous work, which is available within the publication and associated supplementary materials [5]. There were 126 immune measures, from which we removed six as is explained in the next section. Therefore, there are 120 immune measures available for each mouse. These immune measures are classified into six categories, two of which are further divided into sub-categories, as shown in Fig 1.

**Fig 1.**
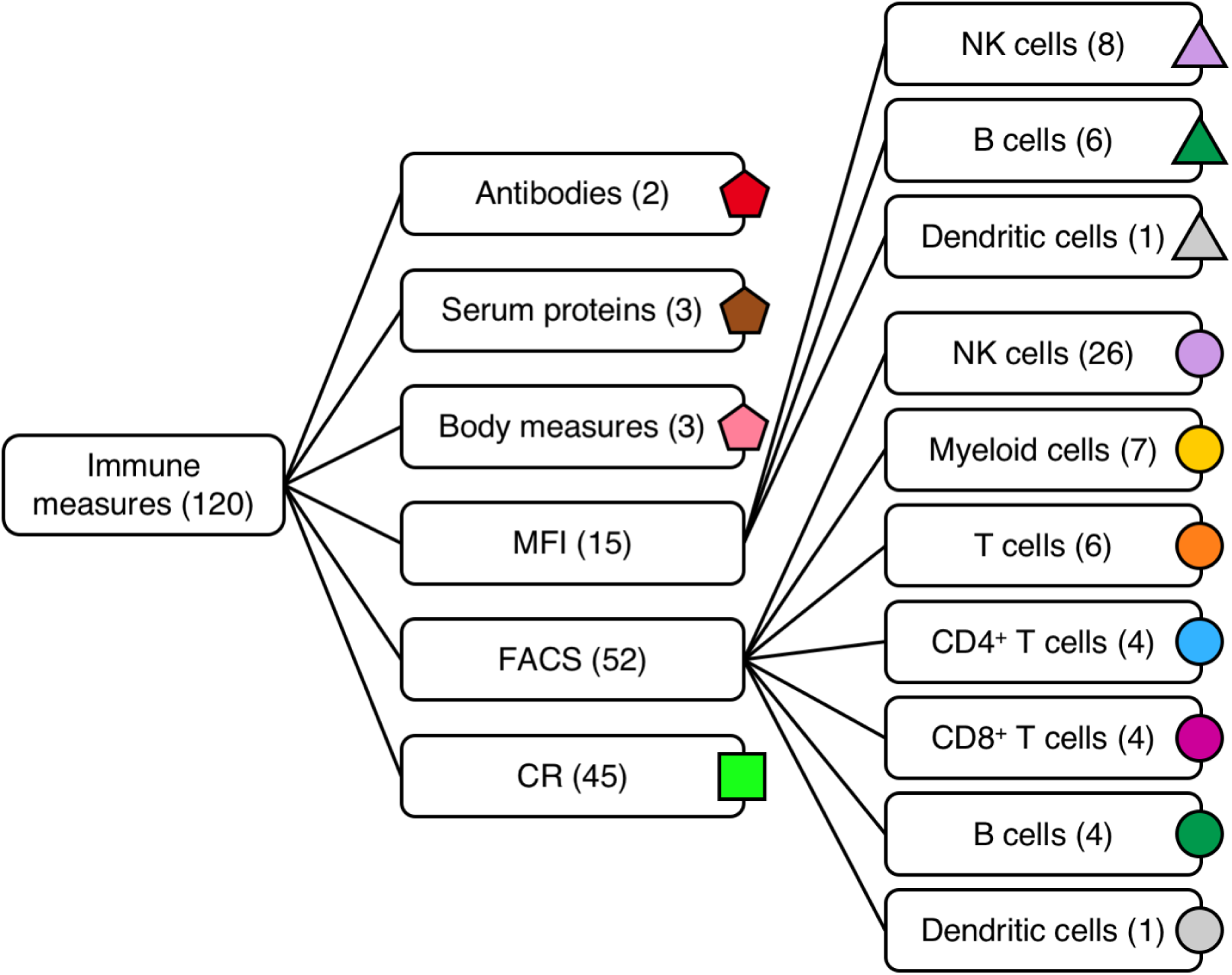
Categories of immune measures. The numbers in the parentheses are the number of immune measures within that category. The symbol and colour associated with each category are used in the subsequent figures. The MFI category and the FACS category are divided into three and seven sub-categories, respectively.

The first category is antibodies, which has two measures: the serum concentration of immunoglobulins G (IgG) and E (IgE). The second category is serum proteins, which has three measures: concentration of the acute phase proteins serum amyloid P (SAP), Haptoglobin, and alpha-1 antitrypsin (AAT). The third category has three non-immunological body phenotype measures: the body mass, spleen mass and the number of spleen cells. The fourth category is *ex vivo* activation state measured as mean fluorescence index (MFI) and has a total of 15 measures. We grouped the 15 measures into three sub-categories: MFI of natural killer (NK) cells (eight measures), MFI of B cells (six measures), and MFI of Dendritic cells (which has one measure). The fifth category is fluorescence activated cell sorted (FACS)-derived percentages of cells and has a total of 52 measures grouped into seven sub-categories: NK cells (26 measures), Myeloid cells (seven measures), T cells (six measures), CD8^+^ T cells (four measures), CD4^+^ T cells (four measures), B cells (four measures) and Dendritic cells (one measure). The sixth category is cytokine responses (CRs). A CR is the concentration of a cytokine produced after *in vitro* stimulation. There are nine types of cytokines: Interleukin (IL)-1*β*, IL-4, IL-6, IL-10, IL-12p40, IL-12p70, IL-13, IFN-*γ* and Macrophage Inflammatory Protein (MIP)-2*α*. Each of the nine types of cytokines is produced after four types of stimulation and one control. Three stimulations are with pathogen-associated molecular patterns (PAMPs), namely CpG, peptidoglycan (PG) and lipopolysaccharide (LPS). The other stimulation is with a mitogen, specifically anti-CD3 and anti-CD28 antibodies (CD3/CD28). The control culture is unstimulated (RPMI). Thus, the nine types of cytokines combined with the five culture conditions gives a total of 45 CR measures.

### Data preprocessing

We preprocessed the data as follows. The concentrations of cytokines were measured using Bioplex Pro kits (M60-009RDPD & MD0-00000EL, Bio-Rad, UK). Therefore, the range of cytokine concentrations for which data were robust is defined by empirically derived standard curves. When the observed concentration of a cytokine fell outside the standard range of these assays [5], the readings were classified as out of range (OOR), being either below (“OOR*<*”) or above (“OOR>”) the standard range. We set “OOR *<*” measures to 0.001 [5], and treated “OOR >” as a missing value.

We removed the age and IgA measures [5] because they were absent for all the laboratory mice. In the FACS category we removed the %D+G2Hi+G2Lo+H+, %D+G2Hi+G2Lo+H-, %D-G2Hi+G2Lo+H+ and %D-G2Hi+G2Lo+H-measures because they were absent for all wild and laboratory mice. As a result of removing these six measures, we had a final total of 120 immune measures.

Many of the 120 immune measures had missing values, i.e., they were not observed for some mice. In particular, in the CR category there were 223 wild mice (48%) and 75 laboratory mice (74%) without any of the 45 CR measures. Because of the central importance of cytokines in immune responses, we decided to remove these mice instead of removing the CR measures. Before removing these mice, the percentage of missing values in the whole data set was 34% for the wild mice and 37% for the laboratory mice. After removing the mice that had all of the CR measures missing, the data set consisted of 237 wild mice with 12% of missing values and 27 laboratory mice with 12% of missing values. In the CR category, the percentage of missing values dropped from 49% to 1.3% for the wild mice and from 74% to 1.5% for the laboratory mice after this removal.

Finally, we imputed the remaining missing values. For each of the immune measures that had missing values, we calculated the average value from the available observations and replaced the missing values by this calculated average. We carried out this imputation procedure separately for the wild and for the laboratory mice.

### Correlation networks

We built correlation networks separately for the wild mice and for the laboratory mice as follows. We used the *N* = 120 immune measures as nodes of the network. We then calculated the Pearson correlation coefficient between each pair of immune measures by regarding the mice as samples. We placed an edge if, and only if, the correlation value exceeded a prescribed threshold. The threshold was the largest possible value with which the remaining network was connected (i.e., when the network consisted of a single connected component).

### Community detection

We combined a stochastic block model (SBM) [7] and a consensus clustering approach to uncover the mesoscopic block structure of the correlation networks. In the microcanonical formulation of the SBM that we used, one minimizes the description length of the network [8], such that one partitions the nodes in the network *𝒢* into *B* blocks to find **b** = {*b*_1_,…, *b*_*N*_}, where *b*_*i*_ ∈ {1, 2,…, *B*} is the group membership of node *i*. The probability of generating the observed network *𝒢* given partition **b** is denoted by *P* (*𝒢*|*θ*, **b**), where *θ* is the set of the additional parameters that control the connectivity between blocks. The probability that the observed network *𝒢* is generated by a partition **b** is given by the following Bayesian posterior probability:

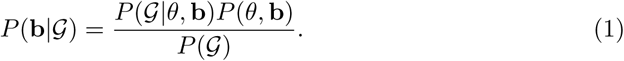

The numerator of Eq (1) can be written as *P* (*𝒢*|*θ*, **b**)*P* (*θ*, **b**) = exp(*-H*), where

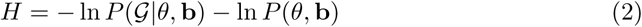

is the description length of network *𝒢*. Maximizing Eq (1) is equivalent to minimizing the description length. We used the non-degree corrected version of the SBM algorithm because it yielded smaller description lengths. Specifically, after 10^3^ minimizing attempts, the average and standard deviation of the description length for the corrected and the non-corrected version was equal to 2575 ± 3 and 2526 ± 4, respectively, for the wild mice and 2118 ± 5 and 2047 ± 5, respectively, for the laboratory mice.

To infer the best partition given by the SBM, we employed a Markov Chain Monte Carlo (MCMC) algorithm [9] using the graph-tool library [10]. We start from an initial partition **b**_**0**_, which is obtained from an agglomerative heuristic method [9]. Then, in each step, we propose a move by selecting a node *i* and choosing a new tentative group membership 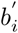 for node *i* with a specific probability that imposes no bias and preserves the ergodicity [9]. The proposed move **b** *→* **b**^*′*^, where 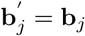 for all *j* ≠ *i*, is accepted or rejected according to the Metropolis-Hastings criterion [11, 12]. This procedure preserves the detailed balance and minimizes the final description length. Each attempt to move a node is applied sequentially, such that nodes are visited one by one in a random order.

When all the *N* nodes have attempted to move once, we say that a sweep has been completed. Because the method is stochastic, a different block structure may appear in each run of the algorithm. To cope with this stochasticity, we implemented the consensus clustering procedure composed of the following three phases. In the first phase, we identify the most probable number of communities of the network. To this end, we ran the algorithm for 10^2^ different initial partitions of the nodes. Then, for each initial condition, we carried out MCMC sweeps and recorded the maximum and minimum values of the description length across sweeps. The node partitions were accepted as being stable if 10^3^ sweeps were completed with less than two record breakings of the description length, where a record breaking is defined by the appearance of the description length value that is larger than the largest value among all the previous sweeps. For each of the 10^2^ initial conditions, we recorded, at an interval of 10^2^ MCMC sweeps, the number of communities 10^4^ times. Then, for each initial condition, we selected the number of communities that appeared the most times in the 10^4^ observations. Finally, we determined the number of communities of the network as the most probable number of communities among the 10^2^ initial conditions. In the second phase, we fixed the number of communities to the value determined in the first phase and inferred the probability that each node belongs to each community as follows. For each of another 10^2^ different initial partitions of nodes, we underwent the same transient period of MCMC sweeps as in the first phase. Then, for each of the 10^2^ initial conditions, we collected 10^5^ configurations at an interval of 10^2^ MCMC sweeps and calculated the fraction of the 10^5^ configurations in which each node belonged to each community. Finally, for each initial condition, we defined the community of each node as the community to which the node belonged the most times. Therefore, at the end of the second phase, we had 10^2^ node partitions. In the third phase, we performed a consensus clustering procedure. To this end, we counted the number of times that each pair of nodes belonged to the same community among the 10^2^ node partitions found in the second phase. We then concluded that any pair of nodes belonged to the same community if they did so in at least 90% of the 10^2^ node partitions. If a node did not belong to the same community with any other node in at least 90% of the 10^2^ node partitions, then this node was judged to form a single-node community.

### Comparing two communities

We estimated the similarity between a single community in the wild mouse network and a single community in the laboratory mouse network in two ways. First, we counted the number of nodes shared by the two communities. Second, we computed the Jaccard index *J* between the two communities, defined by

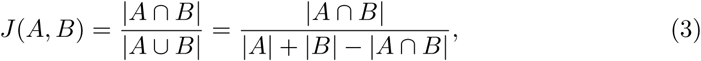

where *A* and *B* are the set of nodes in the two communities being compared, and | *A* |, for example, is the number of nodes in *A*.

### Similarity between community structures

In the statistical analysis, we computed the similarity between two community structures. The networks to be compared were always composed of the same set of nodes but may have different edges. Therefore, the different community structures in the present study are equivalent to different partitions of the same set of nodes. Many similarity measures to compare community structures are available [13]. They can be roughly classified into pair counting methods, cluster matching methods, and information-theoretic methods [13–15]. We used five types of similarity measures, four of which are pair counting methods, with the other one being an information-theoretic method.

The pair counting methods first classify all the possible *N* (*N −* 1)*/*2 pairs of nodes into four categories. We denote by *w*_11_ the number of node pairs that belong to the same community in both networks, by *w*_10_ the number of node pairs that are in the same community in the first network but not in the second network, by *w*_01_ the number of node pairs that are in the same community in the second network but not in the first network and by *w*_00_ the number of node pairs that are not in the same community in either network. The total number of pairs of nodes is given by *w*_11_ + *w*_10_ + *w*_01_ + *w*_00_ = *N* (*N −* 1)*/*2. The four similarity measures based on pair-counting are as follows [13]: the Jaccard index *J* = *w*_11_*/*(*w*_11_ + *w*_10_ + *w*_01_); the Rand similarity coefficien<u>t</u> *R* = 2(*w*_11_ + *w*_00_)*/N* (*N −* 1); the Fowlkes-Mallows similarity coefficient 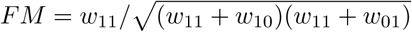; the Minkowski coefficient 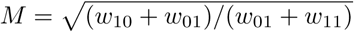.

The other similarity measure based on information theory is the variation of information (VI) [14], which is defined as follows. Denote the two node partitions to be compared by *X* = {*X*_1_, *X*_2_,…, *X*_*k*_} and *Y* ={*Y*_1_, *Y*_2_,…, *Y*_*l*_}, where *k* and *l* are the number of communities in partition *X* and *Y*, respectively. The VI between the two partitions is defined as VI(*X, Y*) = − *Σ*_*i,j*_ *r*_*ij*_[log(*r*_*ij*_*/p*_*i*_) + log(*r*_*ij*_*/q*_*j*_)], where *p*_*i*_ = |*X*_*i*_| */N, q*_*j*_ = |*Y*_*j*_ |*/N* and *r*_*ij*_ = |*X*_*i*_ ∩ *Y*_*j*_|*/N.*

Similarity measures, including the ones described here, strongly depend on the number and size of communities [13]. It is therefore difficult to assert whether the value obtained for each similarity measure is large or small. To circumvent this problem, we calculated the similarity between two partitions relative to that obtained from random partitions. Given a similarity measure *S* calculated from the original data, the Z score is defined as *Z*_*S*_ = (*S − µ*)*/σ*, where *µ* and *σ* are the average and standard deviation, respectively, of the same similarity measure calculated from random partitions. To avoid possible dependence of the Z value on the number of samples (i.e., number of mice) on which the correlation matrix is calculated, we generated random partitions as follows. As an example, for a given similarity measure, consider the comparison between the community structure of the wild male mouse network and that of the wild female mouse network. The male network is derived from the correlation matrix calculated based on the 133 wild male mice; the female network is calculated based on the 104 wild female mice. First, we calculated the similarity measure between the community structure of the male network and that of the female network, giving *S*. Second, we combined males and females and drew 10 pairs of uniformly random networks maintaining the original number of mice in the two networks. In other words, in each randomly drawn pair of networks, one network was derived from 133 mice and the other network was derived from the remaining 104 mice. Third, for each of the randomly drawn pair of networks, we carried out the community detection. Fourth, we calculated the similarity measure between each pair of random networks. Finally, we calculated the Z score on the basis of the 10 random pairs of networks. It should be noted that the Z score of the Minkowski coefficient and the VI was multiplied by −1 because, differently from the other three similarity measures, more similar partitions yield smaller values of Minkowski coefficient or VI. Therefore, a negative Z value indicates that the pair of original networks (e.g., male and female networks) is more different than are pairs of random networks. If the Z score is significantly smaller than zero, we conclude that, in this example, the male and female networks are significantly more different from each other compared to the difference between randomly generated pairs of networks.

We used an interval of 10^2^ MCMC sweeps in the first and second phases of the community detection for the wild and laboratory networks. To obtain the random pairs of networks used to calculate the Z score, and so to statistically compare the wild and laboratory networks, we also used 10^2^ MCMC sweeps. For the other comparisons (i.e., sex, age and geographical site), we used different numbers of MCMC sweeps determined as follows to avoid excessively long computational time. First, for each network to be compared, we discarded transient sweeps before the obtained partitions were sufficiently stable, where the number of MCMC sweeps to be discarded was determined using the same criterion as in the Community Detection section. Then, for each partition we ran a total of *T* = 10^3^ MCMC sweeps. Next, we computed the autocorrelation function *χ*(*t*) of the description length as a function of the number of sweeps *t*, which is given by

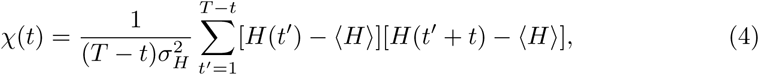

where *H*(*t*) is the description length after *t* sweeps; ⟨ *H*⟩ and *σ*_*H*_ are the average and the standard deviation of the description length, respectively. For each network, we averaged the autocorrelation function of the description length over 10^2^ realizations to obtain a smooth curve of the autocorrelation decay. The time-scale at which the autocorrelation function decays towards zero is called the autocorrelation time *τ*. It is expected that the autocorrelation function declines exponentially at large *t* as *χ*(*t*) *∼ e*^*-t/τ*^ [16]. Therefore, for the other comparisons (i.e., sex, age and geographical site), we set the number of MCMC sweeps as two autocorrelation times *t* = 2*τ*. In fact, 2*τ* for the wild and laboratory networks was equal to 18 and 19 MCMC sweeps, respectively. Thus, using 10^2^ MCMC sweeps guarantees sufficient statistical independence between the configurations drawn.

## Results

### Pairwise correlation between immune measures

Before analysing the data as networks, we compared the raw correlation matrices for the wild and for the laboratory mice. These matrices are shown in Fig 2, where we grouped the immune measures according to the categories of immune measures (Fig 1). Within each category, the immune measures were arranged in descending order of the node strength of the immune measures of the wild mice, where the node strength of an immune measure *i* is defined as the sum of all correlation coefficient values between nodes *i* and *j* over the *N −* 1 possible *j* other nodes.

**Fig 2.**
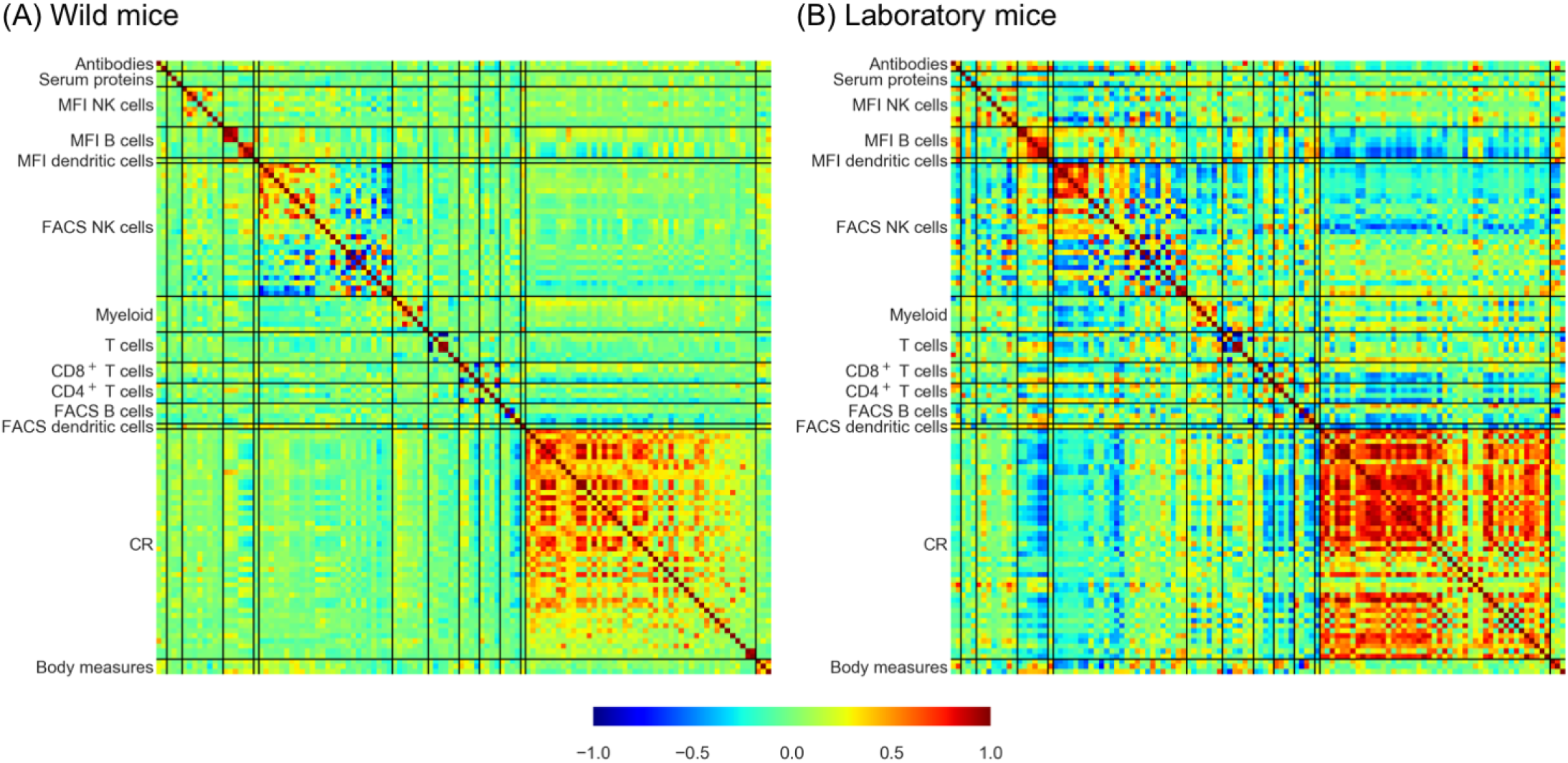
Correlation matrices ordered according to the categories of immune measures. (A) Wild mice. (B) Laboratory mice. The solid black lines separate different categories of immune measure. Within each category, the immune measures are arranged in descending order of the node strength in the wild mouse correlation matrix, using this order for the laboratory mouse correlation matrix too.

Figure 2 shows that, for both wild and laboratory mice, correlations are positive for the majority of the CR pairs. Furthermore, correlations, both positive or negative, tend to be strong within many of the categories of immune measures (i.e., diagonal blocks in Fig 2), for both wild and laboratory mice. In particular, within the FACS NK cells there is a large proportion of immune measure pairs that are strongly, negatively correlated with each other, whereas approximately half of the immune measures in this category are strongly, positively correlated within themselves.

To investigate quantitatively the similarity between the correlation matrices for the wild and laboratory mice, we compared the strength for each node between the wild and laboratory mice, which is shown in Fig 3A. The CR nodes tend to have the largest strength values, while the FACS NK cell nodes tend to have small strength values, which is consistent with the correlation matrices shown in Fig 2. Figure 3A also indicates that the node strength is moderately correlated (with a Pearson correlation value of 0.68; *p <* 10^*-*18^) between the wild and laboratory mice. Comparison of the pairwise correlation values between the wild and laboratory mice shows a moderate correlation (with a Pearson correlation value of 0.51; *p* = 1.8 × 10^*-*17^) (Fig 3B), where each point represents a pair of nodes. Figure 3 suggests that the correlation matrices of wild and laboratory mice show some similarity, but not identity.

**Fig 3.**
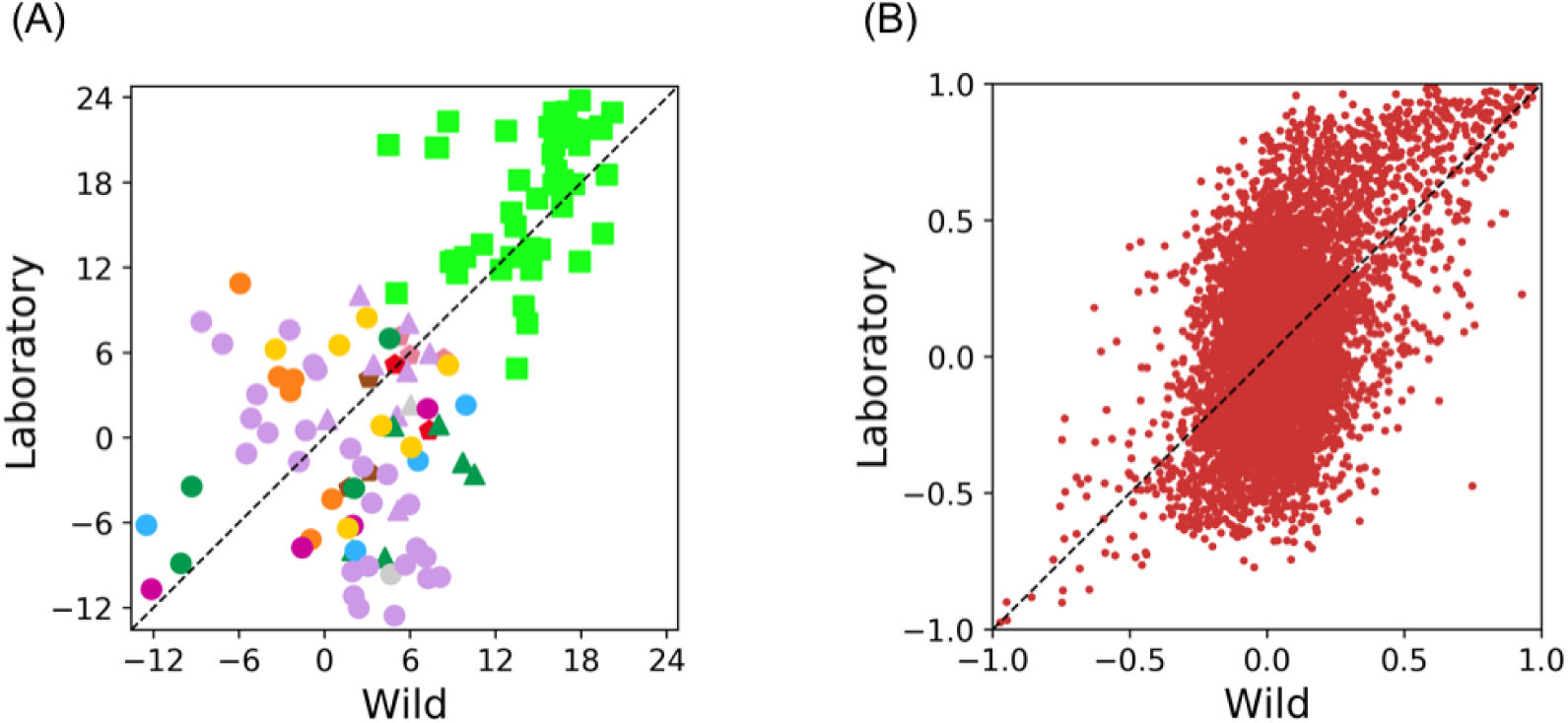
Comparison of the connectivity between the wild and laboratory mice. (A) Strength of individual immune measures. The colour and shape code of the immune measures is as shown in Fig 1. (B) Pairwise correlation coefficients between immune measures. In both, the dotted lines show the diagonal line.

### Community structure of immune measure networks

We next analysed the community structure of networks by thresholding on the pairwise correlation values, considering only positive correlations. We refer to these networks as the wild network and the lab network in the following text. The wild and laboratory mice differ in the distribution of the correlation coefficients among their immune measures and therefore different threshold values were expected in the respective networks. We found that the largest threshold to keep the wild network connected was 0.20, whereas it was 0.42 for the lab network. The edge densities were 0.16 and 0.13 for the wild and lab network, respectively. We confirmed that the following results were reasonably robust when we changed the threshold values to have edge densities of 0.20 and 0.25 for both the wild and lab networks (Figs S1 and S2).

We determined the community structure of the wild and lab networks using the SBM. The final number of communities was equal to seven for both the wild and lab networks, and so the two networks were similar in this respect. In the consensus clustering, the community structure of the wild network is the same for all of the 10^2^ node partitions. The community structure of the networks is shown in Fig 4. Note that community SL in Fig 4D is a single-node community (the FACS NK cell % D+G2Hi-G2Lo+H-) that did not robustly belong to any other community.

**Fig 4.**
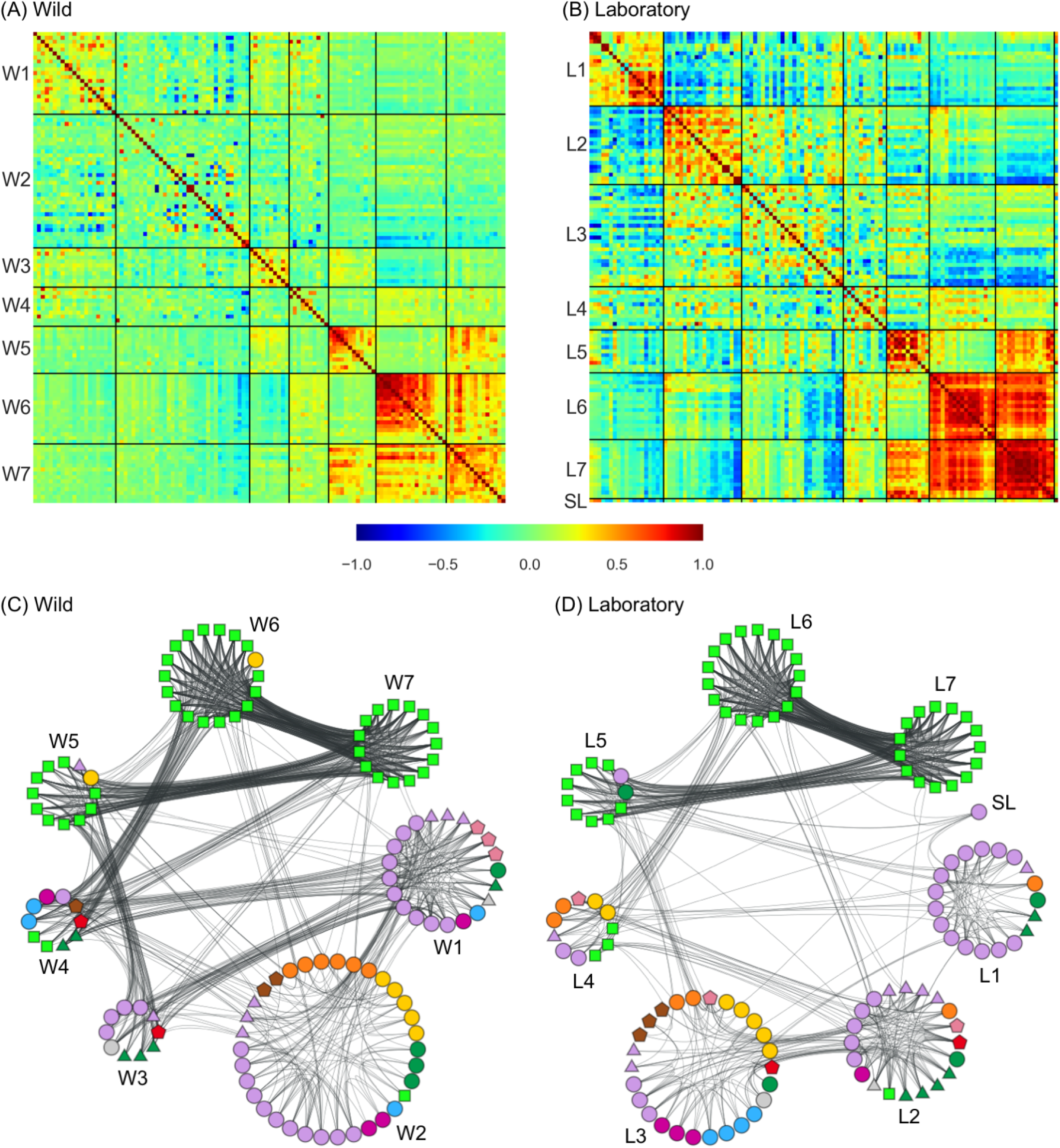
Community structure of the networks. Panels (A) and (B) show the correlation matrix for the wild and laboratory mice, respectively. In each correlation matrix, nodes are ordered according to the block structure detected by the stochastic block modelling. The boundaries between the detected blocks are shown by the solid black lines. The wild and lab networks are shown in (C) and (D), respectively, in a manner respecting the detected block structure.

Both networks have three communities that are almost exclusively composed of CR nodes, which we refer to as the CR communities. Of the 45 CR nodes, 42 and 41 nodes belong to CR communities in the wild and lab networks, respectively. Moreover, the interconnections between the three CR communities is qualitatively the same for both networks. Specifically, there is one CR community that is densely connected to the other two CR communities, which are in turn sparsely connected to each other. In the wild network, community W7 has edge densities of 0.72 and 0.82 with communities W5 and W6, respectively, whereas the edge density between W5 and W6 is equal to 0.06 (Fig 5A). Analogously, in the lab network, L7 has edge densities of 0.60 and 0.83 with L5 and L6, respectively, whereas the edge density between L5 and L6 is equal to 0.03 (Fig 5B).

**Fig 5.**
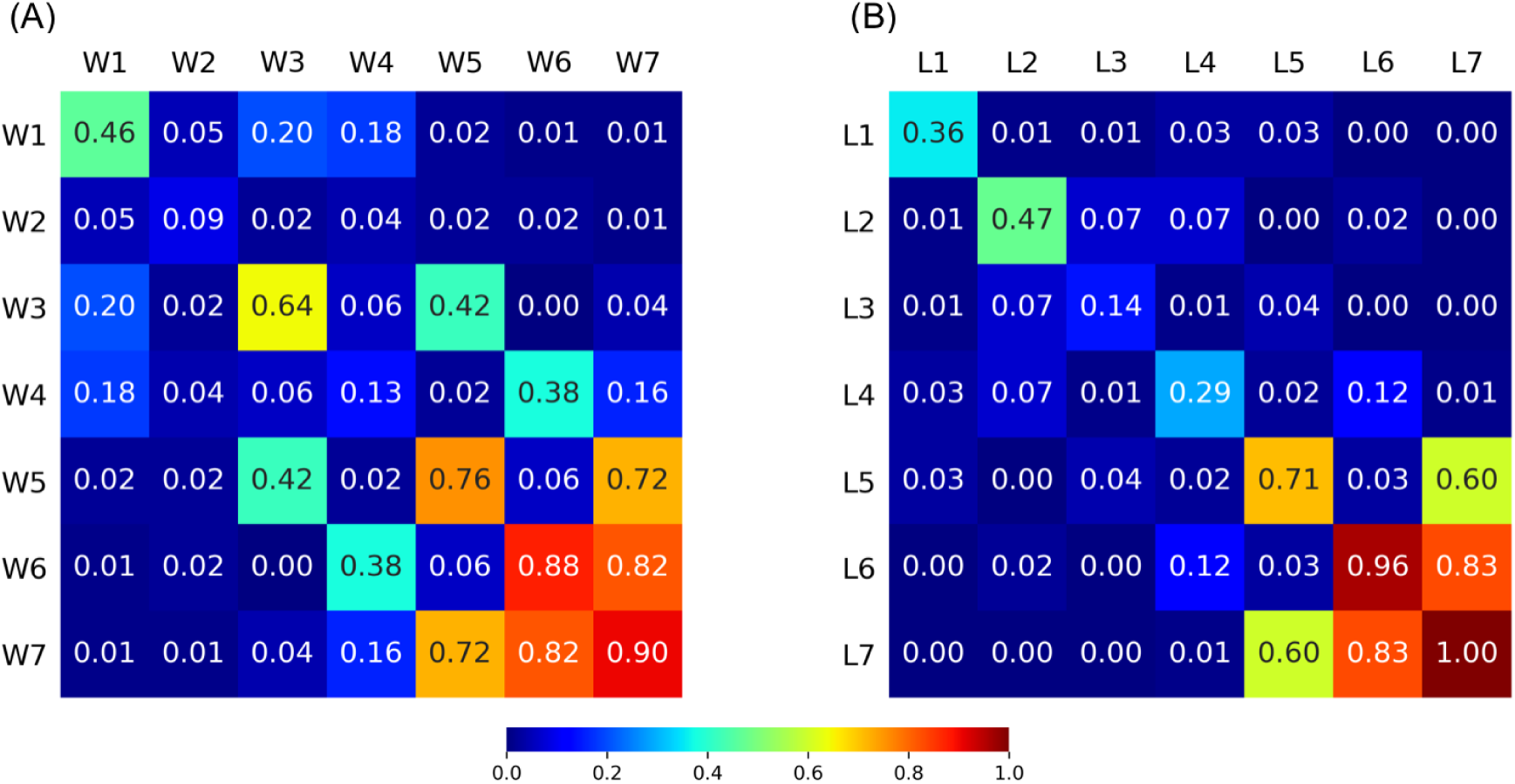
Intra- and inter-community edge densities. (A) Wild network. (B) Lab network.

Despite these structural similarities, the wild and lab CR communities consist of different sets of nodes. We examined this further by computing the number of common nodes and the Jaccard index between each pair of a community in the wild and lab networks (Fig 6). This shows that nodes in the CR communities L5 and L6 of the lab network are scattered among different CR communities in the wild network. Particularly, in the wild network, all of the IL-1*β*, IL-12p70 and IL-13 nodes (with each cytokine type having five nodes) are each within one community, whereas in the lab network only the IL-12p70 nodes are all within one community (Table S1). Moreover, communities W7 and L7, which are strongly connected to the other two CR communities in the wild and lab networks, respectively, have only two CR nodes in common with a Jaccard index of 0.07 (Fig 6). In fact, L7 better corresponds to W6 rather than W7, and W7 better corresponds to L6 than L7. Therefore, although the mesoscale structure of the CR communities is similar between the wild and lab networks, the composition of each CR community is substantially different between the two networks.

**Fig 6.**
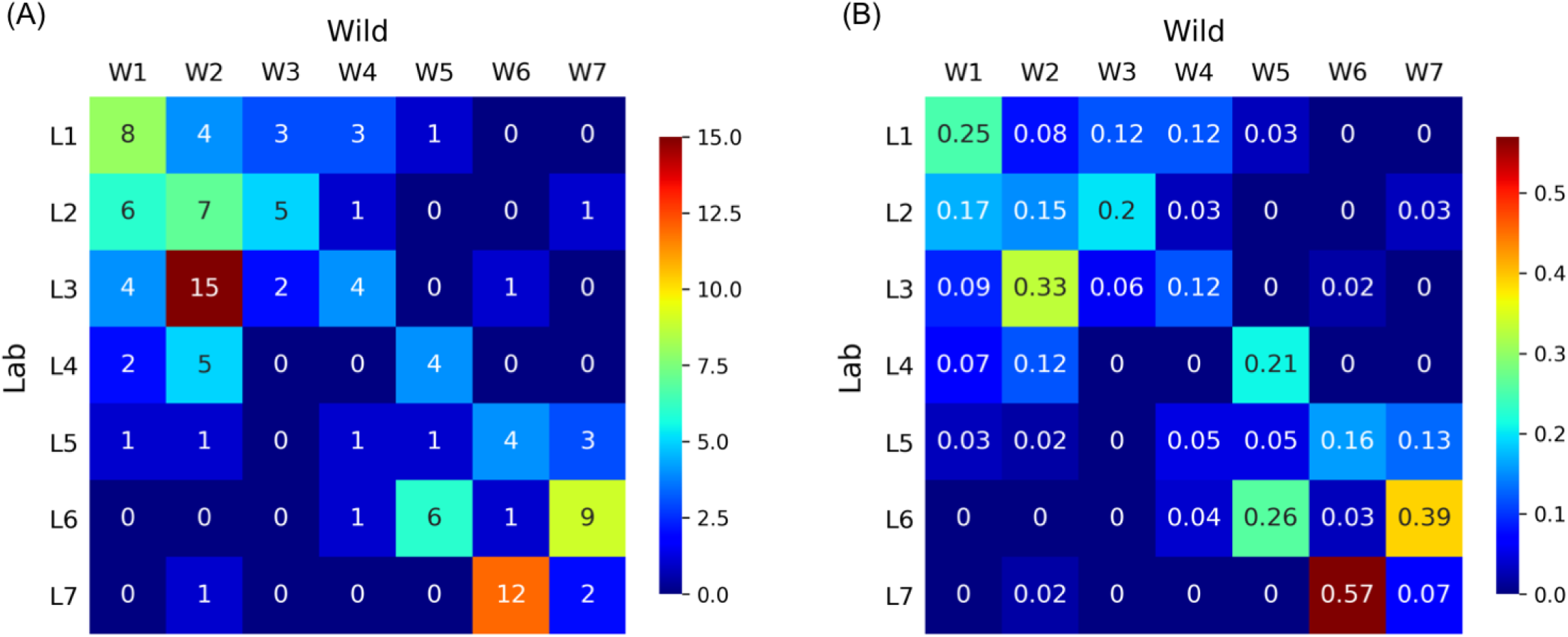
Similarity between communities of the wild and lab networks. (A) Number of common nodes. (B) Jaccard index.

A majority of NK cell nodes (i.e., FACS and MFI nodes) belong to two communities in both the wild network (i.e., communities W1 and W2) and the lab network (i.e., communities L1 and L2). Communities W1 and L1 are predominately composed of NK cell nodes and all the eight nodes that they share are FACS NK cell nodes. Among the seven nodes shared by communities W2 and L2, five are NK cell nodes (three FACS NK cells and two MFI NK cells). In both networks, the edges within each of these two NK cell communities are denser than between the two NK cell communities, which seems to be the main reason why a majority of NK cell nodes are divided between two communities. Consistent with this observation, the pairwise correlations between nodes within L1 and L2 are mostly strongly positive whereas those between L1 and L2 are mostly strongly negative (Fig 4B). This result may imply negative biological regulation among NK cell nodes.

In the wild network, T cells are confined in a single community, i.e., community W2 (Fig 4C), whereas they are dispersed among four communities in the lab network. This result is likely due to greater functional interaction among these cells in wild, antigen-experienced mice, compared with laboratory mice [17].

Communities W2 and L3 are the largest communities in each network and share 15 nodes (Fig 6A). Moreover, these communities contain nodes of different types of immune measures, with this mix being approximately consistent between the wild and lab networks.

We have not considered negative pairwise correlations because the thresholds for creating the networks from the correlation matrices were positive. Mindful that in biological systems negative regulation is common, we also briefly analysed how the immune measures are connected through negative correlations. In these so-called negative networks, node pairs that are strongly negatively correlated were assumed to form the edges. The negative networks have substantially more communities such that their community structure is more difficult to interpret than in the case of the positive networks (Fig S3). However, the CR nodes are predominately grouped together within several communities in the negative networks for both wild and laboratory mice, similar to the situation in the positive networks. In the wild negative network, which had nine communities, 43 out of the 45 CR nodes belonged to one of the two communities that were almost exclusively composed of CR nodes (Fig S3). The lab negative network had 17 communities and 39 CR nodes belonged to one of the four communities predominately composed of CR nodes. Because CR nodes tend to be strongly positively correlated with each other, there is no direct connection between pairs of CR nodes in the negative networks. However, in the SBM, a pair of nodes tend to belong to the same community if they have similar patterns of connectivity to the other nodes in the network. This is the case even if the direct connectivity between the two nodes is not strong, or even absent. This property of the SBM explains why various CR nodes belong to the same community despite the absence of the direct connectivity between them. The results for the negative networks suggest that, for both wild and laboratory mice, the groups of CR nodes have relatively similar patterns of negative connectivity towards other CR or non-CR nodes.

### Comparison of community structures

We next wanted to move from comparing individual nodes and categories of immune measures between wild and lab networks, to statistically comparing the network structure of the wild and lab networks. We also wished to statistically compare networks that address additional features of wild mouse populations that may affect their immune networks, specifically mouse sex, mouse age and the geographical site where the wild mice were collected [5, 17].

In the wild *vs*. lab comparison, there were 237 wild mice and 21 laboratory mice. In the male *vs*. female sex comparison, there were 133 males and 104 females. In the age comparison, we compared 120 old mice, which we defined to be more than 8.5 weeks old, and 117 young mice, which were less than 8.5 weeks old. We selected the age threshold of 8.5 weeks to make the size of the two groups approximately equal in size. In the site comparison, there were 71 mice from the HW site (a mixed arable and beef farm near Bristol, UK) and 166 mice from the other sites [5]. We selected this particular site because it was the single site where most wild mice were sampled.

We compared each pair of networks in terms of the five similarity measures for community structure. For each similarity measure, as a reference, we also randomised the mice to create pairs of networks whose sizes matched those of the original networks. On the basis of the values of the similarity measure for the randomised pairs we calculated the Z score. A negative Z score implies that the pair of original networks are more different than are the pairs of random networks.

The Z score for each of the four comparison types and each of the five similarity measures is shown in Table 1. The largest absolute value of the Z score was obtained when the wild and lab networks were compared with the Rand coefficient. In this case, wild and laboratory community structures are significantly different from each other as compared to pairs of randomised networks (Z = *-*2.93). This result is consistent with those in the previous section, where we have shown that the set of nodes belonging to a community differs between the wild and lab networks, although there is a resemblance in the inter-community connectivity between the wild and lab networks. The Rand coefficient is highly sensitive to the size of communities in the partitions being compared [13]. Both the wild and lab networks have large communities, whereas the randomly generated networks in general do not have similarly large communities. This factor seems to account for this large negative Z score.

**Table 1.** Similarity of the community structure of pairs of networks shown as the Z score. The similarity measures used are the Jaccard index, Rand coefficient, Fowlkes-Mallows coefficient (FM), Minkowski coefficient and the variation of information (VI).

|  | Jaccard | Rand | FM | Minkowski | VI |
| --- | --- | --- | --- | --- | --- |
| Wild *vs.* laboratory | 0.24 | $-2.93$ | 0.16 | 1.34 | 0.19 |
| Male *vs.* female | $-0.37$ | $-0.09$ | $-0.44$ | $-0.48$ | $-1.20$ |
| Young *vs.* old | $-0.96$ | 1.16 | $-1.05$ | $-0.86$ | $-0.89$ |
| HW *vs.* other sites | $-1.42$ | 0.59 | $-1.73$ | $-0.85$ | $-2.04$ |

However, Table 1 shows that, for all but two combinations of the comparison type and similarity measure, the Z scores are below a 95% significance level (i.e., | Z | *<* 1.96). This is even the case for the comparison between the wild and lab networks with the other four similarity measures. For the other three comparisons, we did not find significance except in one case, the geographical comparison with the VI similarity measure with a Z score of *-*2.04. However, when correcting for the multiple comparisons that we have made, neither of these two large negative Z scores remain significant. Therefore, formally we conclude that we cannot detect differences in the immune network structure between wild and lab, young and old, male and female mice, or mice from different geographical sites. However, our results, taken as a whole are strongly suggestive of immune networks differing between laboratory and wild mice, and possibly between mice from different geographical locations too. This general failure to achieve statistical significance may be due to the fact that the number of samples (i.e., mice) from which the correlation matrices were calculated were relatively small as compared to the number of nodes (i.e., immune measures).

## Discussion

We compared the structure of wild and lab mouse correlational networks that were composed of a range of biologically interconnected immune measures. We found that the wild and lab networks were similar in some aspects, such as the strong but chain-like connectivity among the three CR communities, and division of NK cell nodes into two communities. This similarity was not completely unexpected since both wild and laboratory mice are the same species, and so have the same immune system with the same components, which will necessarily have some, even substantial, shared function. However, we also found considerable differences between the two networks, such as the identity of nodes constituting the three CR communities and different arrangement of the T cell nodes. The difference between the wild and lab networks shows how the same immune system can behave differently in animals leading different lives [5].

We found that immune measures of one category can have membership of distinct network communities. In particular, the CR nodes were arranged into three communities in both wild and lab mouse networks, with those communities having similar connectivity; specifically, one community was strongly connected with the other two, but these two other communities were not strongly connected to each other. Despite this mesoscale similarity, the node composition of these communities differs substantially between the wild and lab networks. This result might reflect different regulation of the cytokine network in wild and lab mice (as in [5]), itself a consequence of the different antigenic environment experienced by wild and laboratory mice. The two largest communities in the wild and lab mouse networks (W2 and L3, respectively) both predominantly consist of a mixture of cellular populations. Indeed, these are cellular populations that are known to interact and to cross regulate each other, perhaps most notably the CD4+ and CD8+ T cell populations.

The NK cell nodes were concentrated in two communities in each network – W1 and W2, and L1 and L2 – though these communities also contained other node types. The nodes within W2 contain many (specifically, eleven) markers of naïve, antigen-inexperienced cells (NK, T and B cells), whereas in the lab network these eleven were in L1, L2, L3 and L4. The lab mice are less antigen experienced, compared with the wild mice, and this would therefore appear to be reflected in the different arrangements of the nodes within these communities. These different arrangements of nodes representing these cellular markers may also be because of different immunological states of wild and laboratory mice.

There has previously been some use of network concepts for investigating the immune system and immune responses, but this has not been extensive. A number of different network approaches have been used, which also include frameworks that do not use networks in the sense of graph theory as in the work we present here [18–22]. There are different types of cytokine and immune networks that are based on graph theory and network analysis. First, there are cytokine networks that represent the state transition pathways of cells, in which nodes are cells and edges are cytokines that mediate a state transition of the cell [19, 23, 24]. Second, and perhaps the most common, is where signalling networks of immune systems are analysed, where intercellular signalling proteins (which can include cytokines) are nodes and activation and inhibition of a node by another defines directed edges [21, 25, 26]. A third type are co-expression networks, in which nodes are abundance of cytokines or other types of cells or proteins, and undirected edges represent the co-expression of a pair of immune markers. The strength of association between each pair of nodes has been quantified, for example, by the Pearson correlation coefficient [27, 28], as in the present study, or mutual information [29, 30]. These previous studies analysed data obtained from healthy human donors [28], patients in preterm labour [27], individuals with Chronic Fatigue Syndrome [29] or Gulf War Illness [30]. A final example of network analysis used the proteome of sub-populations of immune cells, clustered these proteins into functional modules, and then used known interactions among proteins to construct networks showing putative interactions within and among functional modules [28]. In the present work, we measured the correlation between a range of different measures of immune state, which included cytokine responses, the number, type and state of different cell populations, and the concentration of serum proteins, in wild and laboratory mice. Therefore, our networks can be regarded as co-expression networks, though the scale of these networks is larger than previous work.

The present study has some limitations. First, we necessarily discarded a considerable portion of data during our pre-processing steps due to missing values. The number of lab mice with sufficient immune measures after pre-processing was small (i.e., 27). However, it is of note that these laboratory animals are genetically inbred and maintained under standard conditions, all aiming to substantially reduce the inter-mouse variability. Among the wild mice, many animals, particularly the young ones, were small, which limited the amount of serum and the number of splenocytes that could be obtained, thereby limiting the amount of immunological data that could be obtained from them. Studies of wild animals will routinely confront such limitations. Therefore, the preprocessing of data that we have undertaken may provide a model for how this matter can be addressed in similar analyses of analogous data of other wild populations. A larger dataset would significantly enhance the precision of the present findings. Second, we did not find consistent patterns in the statistical comparison of the four pairs of groups (wild *vs*. lab; male *vs*. female; young *vs*. old; HW site *vs*. other sites). Rather, the results depended on the similarity measure we used and only a few combinations of the similarity measure and the comparisons that we made yielded significant Z scores. However, adjusting for these multiple comparisons means that none of these results remains formally significant. As above, this may be due to the relatively small size of the data set, compared to the number of nodes. Specifically, to compare two groups of wild mice, such as male and females, we divided the 237 wild mice into two groups. The number of nodes, 120, may be too large for this quantity of data. The same limitation was acknowledged in a study of co-expression immune response networks in which the number of participants in each group was relatively small [30]. However, this is the best that we could achieve with the available data. Sampling wild mice and making these multiple immune measures is a substantial piece of work and we note that this data set is the largest existing immunological data set of wild rodents. Third, we applied a simple thresholding to the correlation matrices to create the correlation networks. The choice of the threshold value was not determined *a priori*, but rather determined functionally by using the highest value that resulted in a single, connected component network [31, 32]. A method to select the threshold value based on an optimisation [32] and the use of network quantities directly applied to correlation matrices [33–35] are alternative options that could be used in the future. We did not opt to use these methods because the optimisation of the trade-off with the wiring cost [32] is not relevant to the present data and stochastic block models directly applicable to correlation matrices are not known.

The network analysis of wild mouse populations presented here is a novel analysis of a novel dataset. It generates a holistic view of the mammalian immune system and the dynamics of its function as the animals themselves interact with diverse and dynamic environments. Humans are, arguably, immunologically more akin to wild mice than to laboratory mice, and the diversity of immune state among people could also be analysed using network analyses. Such an approach may usefully be used to ever better understand the vertebrate immune system and its function in protecting individuals from infection and disease.

## Acknowledgments

The wild and laboratory mouse data set was generated during NERC grant NE/I022892/1. We would like to thank the Jean Golding Institute for seed-corn funding. This study was financed in part by the CoordenaÇão de AperfeiÇoamento de Pessoal de Ní vel Superior - Brasil (CAPES) - Finance Code 001. We thank Eleanor Riley for useful discussions.

